# Female reproductive fluid composition differs based on mating system in *Peromyscus* mice

**DOI:** 10.1101/2022.03.18.484937

**Authors:** Kristin A. Hook, Catherine Liu, Katherine A. Joyner, Gregg A. Duncan, Heidi S. Fisher

## Abstract

Post-copulatory sexual selection is theorized to favor female traits that allow them to control sperm use and fertilization, leading to the prediction that female reproductive traits that influence sperm migration should differ between polyandrous and monogamous species. Here we exploit natural variation in the female mating strategies of closely related *Peromyscus* mice to compare female traits that influence sperm motility – the viscosity, pH, and calcium concentration of fluids in the reproductive tract – between polyandrous and monogamous species. We find that the viscosity and pH, but not calcium concentration, of fluids collected from both the uterus and the oviduct significantly differ between species based on mating system. Our results demonstrate the existence of a viscosity gradient within the female reproductive tract that increases in monogamous species but decreases in polyandrous species. Both species have a more alkaline environment in the uterus than oviduct, but only in the polyandrous species did we observe a decrease in calcium in the distal end of the tract. These results suggest that fluid viscosity and pH in the female reproductive tracts of these mice may be influenced by post-copulatory sexual selection and provide a promising potential mechanism for female sperm control given their importance in modulating sperm behavior.

## INTRODUCTION

When females mate with more than one male within a single reproductive cycle (i.e., polyandry), post-copulatory sexual selection is hypothesized to favor male reproductive traits that allow them to outcompete rivals in their race to fertilize a females’ ova (i.e., sperm competition; Parker 1970) and female traits that allow them to preferentially bias sperm use in favor of certain males over others (i.e., cryptic female choice; Thornhill 1983; Eberhard 1996) and to prevent polyspermy (Kim et al. 1996; Firman and Simmons 2013; Firman 2018). Consequently, female reproductive traits that enable post-copulatory control and selective fertilization of ova are predicted to occur in polyandrous – but not monogamous – species. This prediction has been poorly tested, however, likely due to lack of attention to female reproductive traits compared to male reproductive traits in post-copulatory sexual selection studies (Orr et al. 2020), the technical challenges of examining covert mechanisms of ‘cryptic female choice’ for internally fertilizing species (reviewed in Firman et al. 2017; Ng et al. 2018), and the difficulty of disentangling these female mechanisms from sperm competition and sexual conflict (Simmons and Wedell 2020).

Despite the challenges with demonstrating cryptic female choice, several studies across diverse taxa have provided empirical support that females are capable of preferentially biasing and controlling sperm use (reviewed in Firman et al. 2017; Firman 2020). For example, in feral fowl (*Gallus domesticus*), females use muscular contractions to eject sperm from socially subdominant males to prevent insemination and fertilization of their ova (Pizzari and Birkhead 2000); in Japanese macaques (*Macaca fuscata*), females increase orgasm-like muscular contractions after mating with a socially dominant male (Troisi and Carosi 1998), which increases sperm retention within their reproductive tract (Baker and Bellis 1993); and in red flour beetles (*Tribolium castaneum*), females appear to be in control of the observed sperm precedence patterns based on male copulatory behavior (Edvardsson and Göran 2000). There is also evidence that physical structures within the female reproductive tract enables female control of sperm use across taxa. For instance, in fruit flies (*Drosophila melanogaster*), female sperm storage organs allow them to control the timing and use of sperm stored after copulation with multiple males (Manier et al. 2010). Moreover, many female vertebrates possess a tube-like passageway to their ovaries (i.e., the oviduct), and there is evidence in birds and mammals that features of this structure (Holt and Fazeli 2016), such as its length (Gomendio and Roldan 1993; Anderson et al. 2006), positively correlate with relative testis size, a proxy for sperm competition level (reviewed in Simmons and Fitzpatrick 2012; Vahed and Parker 2012; Lüpold et al. 2020). These findings suggest that the variable structural architecture of the female reproductive tract may have evolved to regulate sperm uptake (Suarez 2008; Tung and Suarez 2021) by selecting for only those sperm cells that are able to bypass its challenging features (Holt and Fazeli 2016; Suarez 2016) while excluding pathogens or microbes (Tung et al. 2015; Holt and Fazeli 2016; Rowe et al. 2020).

The composition of fluids within the female reproductive tract may provide yet another potential mechanism of female control within internally fertilizing species, given that their biochemical properties have been shown to change after insemination, vary throughout the tract, and modulate sperm motility and migration to the ova and, thus, the outcomes of fertilization (reviewed in Holm and Ridderstråale 1998; Hunter et al. 2011; Kirkman-Brown and Smith 2011; Holt and Fazeli 2016; Ng et al. 2018; Gasparini et al. 2020). For example, fluids within the reproductive tract can vary in their viscoelastic properties (Johansson et al. 2000; Rodríguez-Martínez et al. 2005; Suarez 2016), which can influence sperm motility patterns and trajectory (Tung et al. 2015; Holt and Fazeli 2016; Tung and Suarez 2021). In humans, mucus coats the entire female reproductive tract, and sperm must swim through viscoelastic cervical mucous as well as the cumulus mass en route to the oocyte (Kirkman-Brown and Smith 2011); a previous study used artificial insemination to demonstrate that this change in fluidic properties effectively serves as a barrier, allowing only more motile and morphologically normal sperm to pass through to the oviduct (Hanson and Overstreet 1981). Moreover, a pH gradient has been demonstrated throughout the female reproductive tract of different mammals, with the uterine environment being more acidic (i.e., less alkaline) than the oviductal environment (reviewed in Ng et al. 2018). Alkaline environments have been shown to increase sperm velocity and induce sperm hyperactivation in mammals, in part through the activation of essential sperm-specific CatSper protein channels (Kirichok et al. 2006; Lishko et al. 2010) that increases sperm intracellular calcium concentrations (Ho and Suarez 2001; Suarez 2008) and subsequently increases their flagella beat frequency and velocity (Brokaw et al. 1974; Suarez et al. 1993). In boars (*Sus scrofa*), high calcium environments lead to greater sperm motility, whereas low calcium environments cause sperm cells to stick to oviductal epithelium and be less motile (Petrunkina et al. 2001). Together these studies suggest that the chemical composition of female reproductive fluids provides a promising mechanism for female sperm control driven by post-copulatory sexual selection, but whether these fluidic properties differ between polyandrous and monogamous species remains unknown.

In this study, we test whether the composition of female reproductive tract fluids diverge between species that have evolved under divergent mating systems in *Peromyscus* mice. More specifically, we collected fluids from two distinct regions of the reproductive tract – the uterus and the oviduct – for three polyandrous species (*P. maniculatus, P. leucopus*, and *P. gossypinus*) and their closely related monogamous congeners (*P. californicus, P. eremicus, and P. polionotus*; Turner et al. 2010; Bedford and Hoekstra 2015). From these fluids, we measured viscosity, pH, and calcium concentration, all of which have been shown to significantly impact sperm movements in other taxa. We compared these physiological properties between species that evolved under polyandry to those that evolved under monogamy to examine associations between mating strategy and potential mechanisms of post-copulatory female control. From these data, we were also able to establish for each species whether a gradient for each of these properties exists within the reproductive tract and indirectly assess how that might impact sperm motility and their unique ability to form collective groups within these mice (Hook et al. 2022).

## MATERIALS AND METHODS

### Female fluid collection

We obtained captive *Peromyscus maniculatus bairdii, P. polionotus subgriseus, P. leucopus, P. eremicus*, and *P. californicus insignis* from the Peromyscus Genetic Stock Center at the University of South Carolina, and *P. gossypinus* from Dr. Hopi Hoekstra at Harvard University. We housed the mice in same-sex cages at 22ºC on a 16L:8D cycle in accordance with guidelines established by the Institutional Animal Care and Use Committee at the University of Maryland (protocol # R-Jul-18-38). We sought samples from all available captive *Peromyscus* species and avoided wild-caught specimens to control for variation due to age, nutrition, and sexual experience. We collected fluid from females in estrus (identified by methods in Byers et al., 2012) to reduce variation associated with the estrous cycle (Hunter et al. 2011; Simons and Olson 2018). We euthanized all focal females via isoflurane overdose and cervical dislocation prior to removing the reproductive tract for fluid extraction.

### Collecting fluid and particle tracking microrheology (PTM)

To collect fluid for viscosity measurements, we removed the female reproductive tract and submersed it in mineral oil at 4ºC until fluid extraction, which took place within 24 hours. To visually distinguish the mineral oil from biological fluid, we dyed the oil blue using a colored gel dye (Wilton Candy Colors, USA) in a 1:75 dye:oil ratio. Under 0.63x magnification (Zeiss Stemi 508, USA), we trimmed the fat surrounding the reproductive tract, unraveled the coiled oviducts, severed the uterus from the oviduct at the utero-tubal junction (UTJ), divided the oviduct in half to separate the lower and upper regions, and submersed in dyed mineral oil. We used glass Pasteur pipettes bent into a u-shape under a flame to push down from one end of the tissue to the other to squeeze fluid within the tissue out into the oil, then collected and centrifuged the samples, removed the oil supernatant and stored fluids at -80ºC (Yuana et al. 2015; Patczai et al. 2017).

To obtain sufficient volume for downstream methods, we pooled the samples for each region from at least ten individuals per species, then warmed pools to 37ºC to simulate natural physiological values, and again centrifuged at 3000 rpm for 3 min to ensure full separation of the mineral oil from the reproductive fluid. We then combined 2µL of fluid with 0.5µL of ∼0.002% w/v suspension of fluorescent nanoparticles (PEG-coated polystyrene particles, PS-PEG). PS-PEG were prepared by coating red fluorescent carboxylate-modified PS spheres (PS-COOH), 500 nm in diameter (ThermoFisher FluoSpheres Carboxylate-Modified Microspheres, 0.5 µm, red fluorescent (580ex/605em), 2% solids, USA) with 5-kDa methoxy-PEG-amine (Creative PEG-Works, USA) via NHS-ester chemistry as previously described (Joyner et al. 2019). We gently reverse-pipetted the mixture to make sure the nanoparticles were homogeneously scattered throughout and pipetted 2.0µL into a 1-mm ID Viton O-ring microscopy chamber (McMaster Carr, USA) and covered with a small circular glass coverslip, both of which were sealed with vacuum grease (Dow Corning, USA) to prevent fluid flow and evaporation, and equilibrated for 30 minutes prior to imaging to reduce dynamic error.

To measure viscosity, we recorded a minimum of three videos of the suspended fluorescent nanoparticles within each fluid sample at a frame rate of 33.33 Hz for 300 frames (10 sec) using an EMCCD camera (Axiocam 702; Zeiss, Germany) attached to an Zeiss 800 LSM inverted microscope and x63/1.20 W Korr UV VIS IR water-immersion objective with image resolution of 0.093μ per pixel. To avoid edge effects on nanoparticle movement, we randomly selected central locations within the chamber for our video recordings. All samples remained at 37ºC during imaging using a stage incubator (PM 2000 Rapid Balanced Temperature, PeCon, Germany). To track the diffusion of PS-PEG nanoparticles in each sample, we used particle tracking data analysis using automated software custom-written in MATLAB (Mathworks, USA). Based on a previously developed algorithm (Crocker and Grier 1996), the program determined the *x* and *y* positions of nanoparticle centers based on an intensity threshold and then constructed particle trajectories by connecting particle centers between sequential images given an input maximum moving distance between frames. Finally, the program calculated the time-averaged mean squared displacement [MSD (**τ**)] as

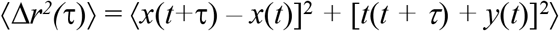

where **τ** is the time lag between frames and angle brackets denote the average over the time points. The MSD of PS-PEG nanoparticles is directly proportional the viscosity of the surrounding fluid. A fast-moving particle (high MSD) reflects a low viscosity fluid whereas a slow-moving particle (low MSD) reflects a high viscosity fluid. Using the generalized Stokes–Einstein relation, measured MSD values were used to compute viscoelastic properties of the hydrogels (Joyner et al. 2020). The Laplace transform of ⟨Δ*r*^2^(*τ*)⟩, ⟨Δ*r*^2^(*s*)⟩, is related to viscoelastic spectrum *G*(*s*) using the equation *G*(*s*) = 2*k*_B_*T*/[π*as*⟨Δ*r*^2^(*s*) ⟩], where *k*_B_*T* is the thermal energy, *a* is the particle radius, *s* is the complex Laplace frequency. The complex modulus can be calculated as *G*\*(*ω*) = *G*′(*ω*) + *G*′′(*iω*), with *iω* being substituted for *s*, where *i* is a complex number and *ω* is frequency. We pooled these data across all particles to characterize female reproductive fluid viscosity per species. Due to technical difficulties, we were unable to collect viscosity data from the upper oviduct for two of the polyandrous species (*P. maniculatus* and *P. gossypinus*). For this reason, we combined viscosity data for both the lower and upper oviducts into a single measure for every focal species.

### Collecting and measuring fluid pH and calcium

Due to the limited quantity of fluids collected from each female reproductive tract for our viscosity measurements, we were not able to also measure pH and calcium using these same individuals. Because we were further limited by the number of available animals in our lab colony, we focused on two species in which ten females were available for this study – the polyandrous *P. maniculatus* and its monogamous sister-species *P. polionotus*. To extract reproductive fluid for these measurements, we dissected the reproductive tract, reserved the right side of each tract for the pH measurements and submersed the left side of each tract into phosphate buffer solution (PBS) at 4ºC for the calcium measurements.

To measure the pH of reproductive fluids, we placed pH strips (Hydrion [9400] Spectral 5.0-9.0) on microscope slides (VWR Plastic Microslides, USA) on a 37ºC warmer (Fisherbrand Isotemp Digital Dry Bath/Block Heater, USA) and under 0.63X magnification (Zeiss Stemi 2000, USA). We trimmed the fat, unraveled the coiled oviduct, placed the elongated tract on a pH strip, and severed the uterus from the oviduct at the UTJ. We then placed the oviduct on another pH strip, cut it in half, pinched the open end of the lower oviduct closed with forceps, and placed it on a third pH strip. We used separate u-shaped glass pipettes to push from one end of each region of the reproductive tract to the other end to squeeze fluid out onto the pH strip to measure pH immediately after release from the tissue (Yeung et al. 2004).

To measure calcium in reproductive fluids, we used a calcium assay kit (Abcam Calcium Assay Kit ab102505, USA). After removing the tissue from PBS, we severed the uterus from the oviduct at the UTJ, cut the oviduct in half to separate the lower and upper oviducts, placed each tissue region in a separate tube containing calcium assay buffer, and ground the tissues with disposable plastic pestles (ThermoFisher, USA). We separated the tissue from the buffer and reproductive fluid by centrifuging at 4ºC for 5 min at 3500 rpm (Eppendorf Centrifuge 5702RH, USA), pipetted the supernatant from each tract region into microplate wells (CellStar 96 Well Cell Culture Plate, USA), added the chromogenic reagent, calcium assay buffer, and calcium standard, and immediately measured the mixture’s absorbance at 575nm using a microplate reader (Thermo Scientific Multiskan FC, USA).

### Statistical analyses

We performed all statistical analyses using R version 3.4.2 (R Core Team 2016) and created all figures using the ‘ggplot2’ package with R (Wickham 2016). We visually inspected all diagnostic plots (qqplots and plots of the distribution of the residuals against fitted values) to validate model normality. In cases where model assumptions of normality were not, we assessed the normality of the response variable using a Shapiro-Wilk test and transformed variables as needed until normality of models was met. Only the best fitting models are reported here.

To assess differences in the viscosity of female reproductive fluids between polyandrous and monogamous species, we used our dataset of individually tracked particles and pooled these data across species with shared mating systems (categorized as either monogamous or polyandrous). We excluded data that were ±2SD of the mean within each sample because they represented clear outliers. All viscosity values were log-transformed to improve model assumptions of normality. We assessed differences in the viscosity of female reproductive fluids in the uterus and in the oviduct using separate linear models (LM), with log viscosity as the response variable and mating system as the predictor variable. We also analyzed fluidic differences in each tract region within each species using separate linear models, with log viscosity as the response variable and the region of the female reproductive tract as the predictor variable. Last, we used the data sets from three focal samples (*P. eremicus, P. gossypinus*, and *P. polionotus*) to statistically compare viscosity measurements to their dyed mineral oil controls using separate linear models, with log viscosity as the response variable and the sample type as the predictor variable. Post-hoc pairwise comparisons were made using Tukey HSD adjustments for multiple comparisons using the ‘LSmeans’ R package (Lenth 2016).

To assess differences in the pH and calcium levels of female reproductive fluids between polyandrous and monogamous species, we used separate linear models for each region of the tract, with either pH or the log calcium measurements included as the response variable and mating system included as the explanatory variable. Last, we used paired t-tests to conduct pairwise comparisons for pH and calcium measurements from each region of the tract within each species to control for differences among individual females.

## RESULTS

All fluid viscosity measures collected from the female reproductive tracts of each focal *Peromyscus* species are reported in Table 1. We found that the viscosity of fluids in both the uterus and the oviduct significantly differed based on mating system in *Peromyscus* mice. More specifically, polyandrous species have significantly more viscous fluid in the uterus (LM: F_1,1255_ = 27.09, p < 0.001) but significantly less viscous fluid in the oviduct (LM: F_1,1755_ = 24.7, p < 0.001) than monogamous species (Figure 1). Within-species analyses revealed that fluids were significantly more viscous when collected from the uterus compared to the oviduct in *P. maniculatus*, P. *leucopus*, and *P. californicus*, but the opposite was true in *P. eremicus* and *P. polionotus*; no difference was observed in the viscosity of uterine fluid or oviductal fluid in *P. gossypinus*, however (Table 1).

**Table 1.** Mean (± SE) viscosity (Pa*s) of female reproductive tract fluids extracted from 5 *Peromyscus* mice that naturally vary by mating system

| Mating system | *Peromyscus* species | Viscosity (Pa * s) |  | Linear model results from within species comparisons |
| --- | --- | --- | --- | --- |
|  |  | Uterus | Oviduct |  |
| Polyandrous | *P. maniculatus* | 0.04 ± 0.02<br>(337) | 0.02 ± 0.01<br>(424) | $F_{1,759} = 175.9$, $p < 0.001$ |
| | *P. leucopus* | 8.8 ± 6.6<br>(115) | 0.006 ± 0.002<br>(17) | $F_{1,130} = 4.528$, $p = 0.035$ |
| | *P. gossypinus* | 0.002 ± 2.2e-04<br>(123) | 0.001 ± 5.8e-05<br>(130) | $F_{1,251} = 0.28$, $p = 0.597$ |
|  | **Mean ± SE** | **1.79 ± 1.32** | **0.01 ± 0.01** |  |
| Monogamous | *P. californicus* | 0.29 ± 0.14<br>(258) | 0.04 ± 0.03<br>(72) | $F_{1,328} = 16.76$, $p < 0.001$ |
| | *P. eremicus* | 0.004 ± 8.1e-04<br>(35) | 0.25 ± 0.25<br>(251) | $F_{1,284} = 28.34$, $p < 0.001$ |
| | *P. polionotus* | 0.35 ± 0.22<br>(389) | 0.86 ± 0.49<br>(863) | $F_{1,1250} = 111.3$, $p < 0.001$ |
|  | **Mean ± SE** | **0.31 ± 0.14** | **0.68 ± 0.36** |  |

The number of tracked particles for each sample is indicated within parentheses. Linear models included log-transformed viscosity values as the response variable and the region of the reproductive tract as the explanatory variable within each species.

We found that the reproductive fluid pH of the focal polyandrous species, *P. maniculatus*, was significantly higher within the uterus (LM: F_1,18_ = 17.05 *p* < 0.001) and oviduct (LM: F_1,18_ = 14.39, *p* < 0.01) compared to its monogamous congener, *P. polionotus* (Table 2, Figure 2). However, we found no differences in calcium concentrations between these species in either the fluids collected from the uterus (LM: F_1,18_ = 2.005, *p* = 0.174) or from the oviduct (LM: F_1,18_ = 0.3423, *p* = 0.566; Table 2, Figure 3).

**Figure 1.**
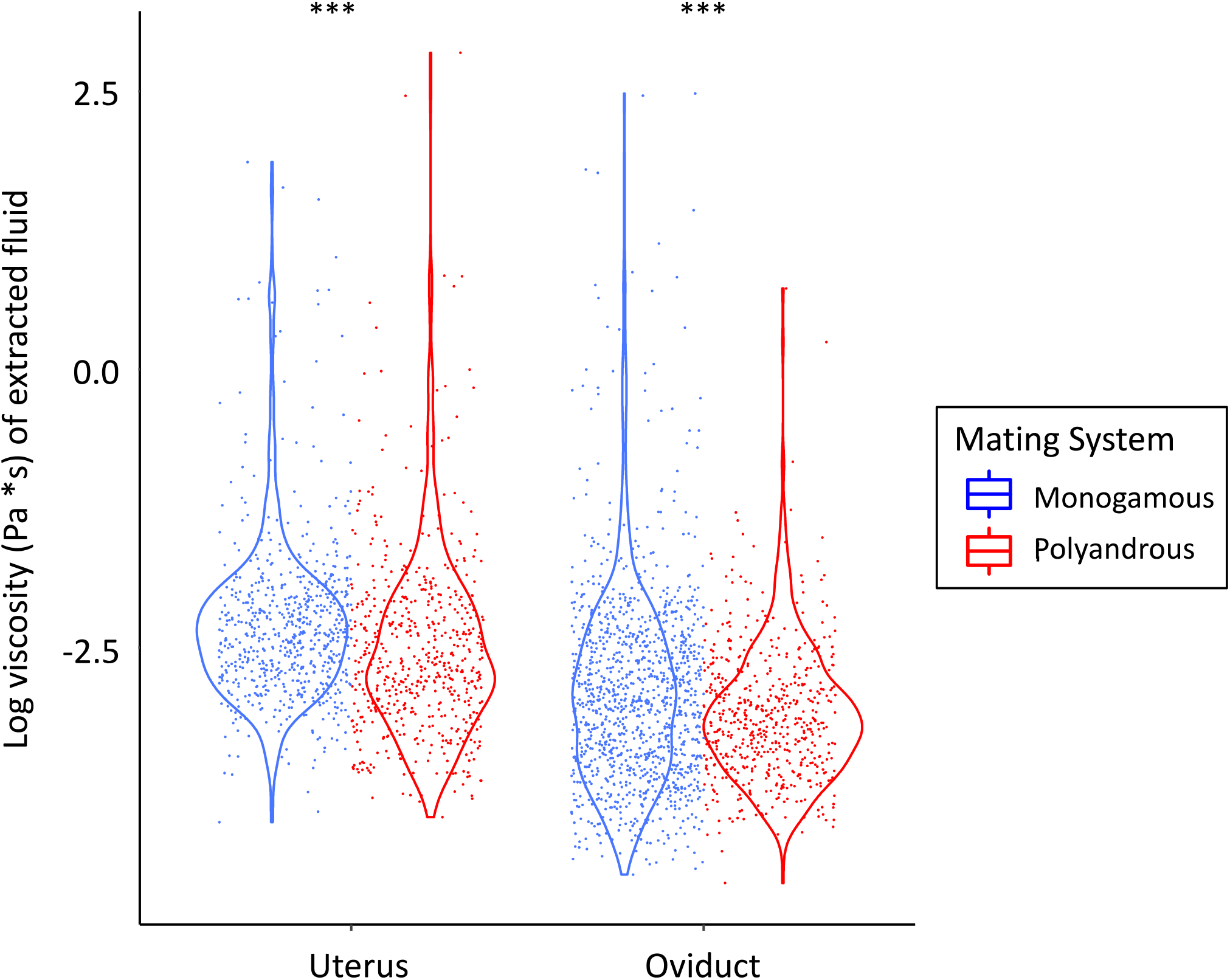
Violin plots of the log-transformed viscosity measurements obtained from fluids collected from two regions of the female reproductive tract within *Peromyscus* mice species with naturally varying mating systems. Blue dots represent particle tracking data obtained from three monogamous species (*P. californicus, P. eremicus*, and *P. polionotus*), red dots represent particle tracking data obtained from three polyandrous species (*P. maniculatus, P. leucopus*, and *P. gossypinus*). Asterisks denote differences within each reproductive region between the mating systems (*** *P* < 0.001).

**Table 2.** Mean (±SE) pH and calcium concentration of female reproductive fluids collected from two sister species of *Peromyscus* mice that vary in their mating system

| Mating system | Species | pH | | Calcium concentration ($\mu\text{m}/\mu\text{L}$) | |
| --- | --- | --- | --- | --- | --- |
|  |  | Uterus | Oviduct | Uterus | Oviduct |
| Polyandrous | *P. maniculatus* | $8.00 \pm 0.00$ | $7.44 \pm 0.03$ | $5.73 \pm 0.97$ | $2.84 \pm 0.42$ |
| Monogamous | *P. polionotus* | $7.76 \pm 0.06$ | $7.28 \pm 0.03$ | $4.11 \pm 0.40$ | $3.22 \pm 0.53$ |

**Figure 2.**
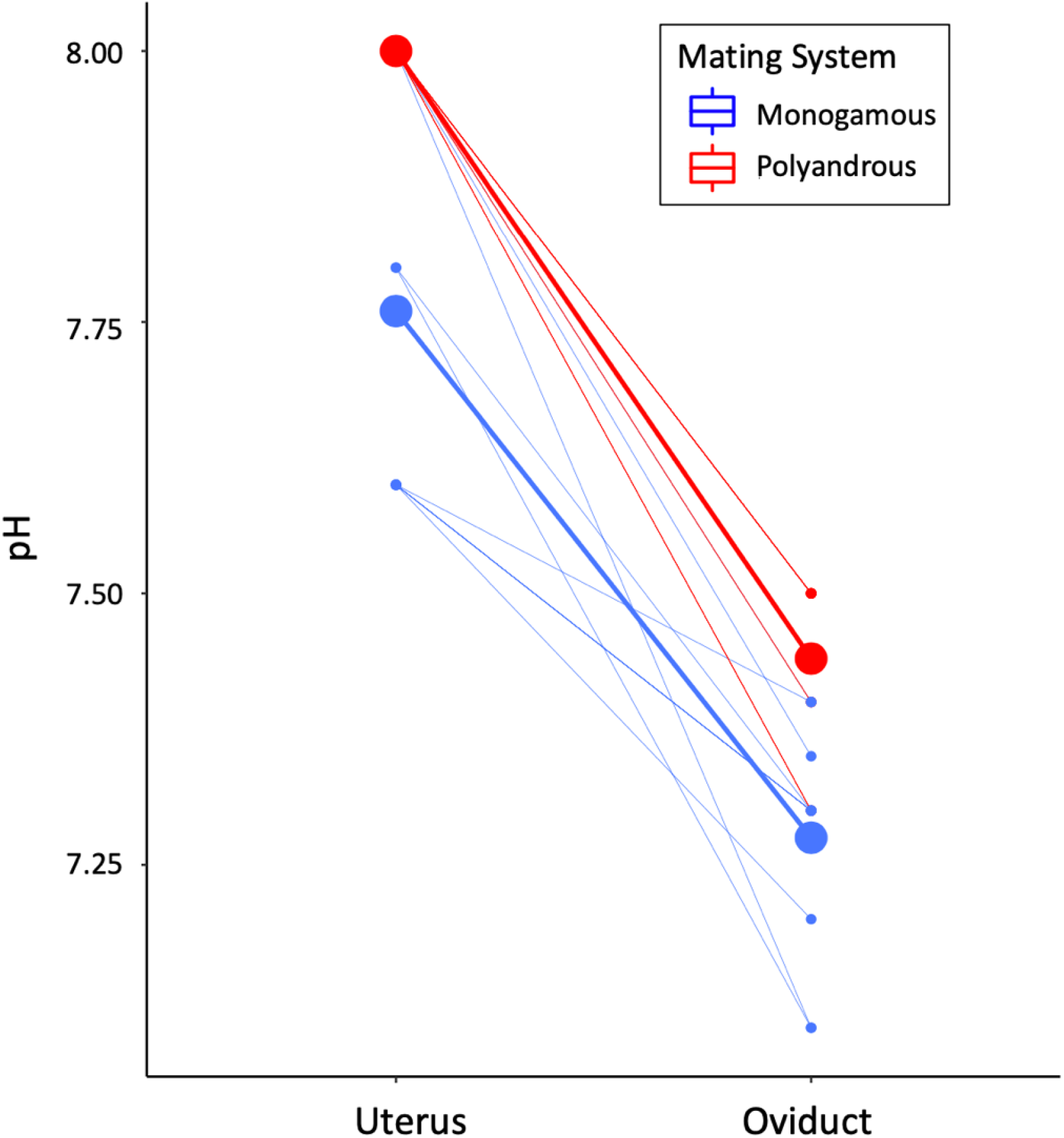
pH measurements obtained from fluids collected from the female reproductive tracts of *Peromyscus* mice species with naturally varying mating systems. In both the uterus and the oviduct, reproductive fluids have a significantly greater pH in the polyandrous *P. maniculatus* (red dots) than its monogamous sister species, *P. polionotus* (blue dots). Small dots denote values measured for individual females; large dots represent the mean values for each species.

**Figure 3.**
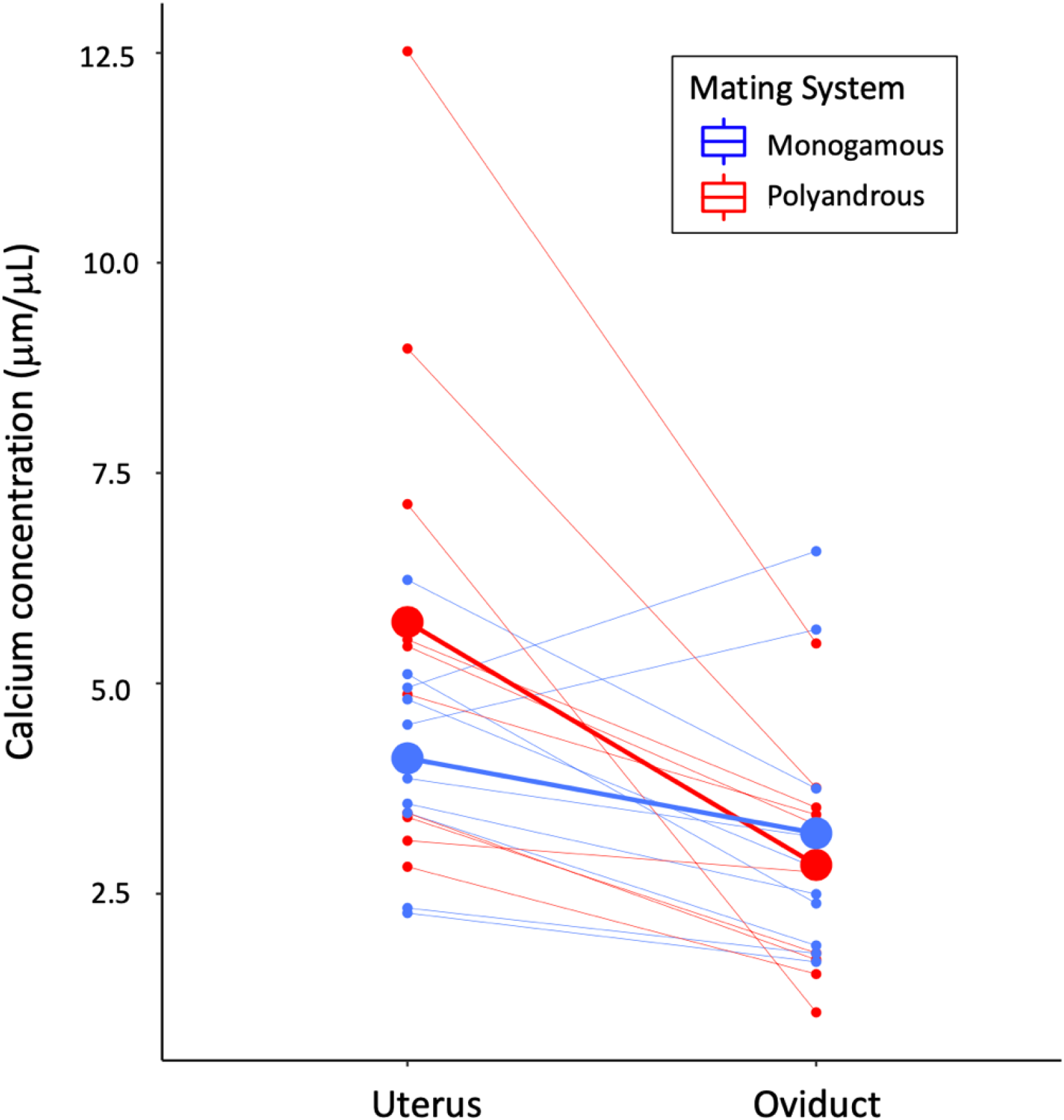
Calcium concentration measurements obtained from fluids collected from the female reproductive tracts of *Peromyscus* mice species with naturally varying mating systems. The calcemic content of reproductive fluids collected from the uterus and oviduct did not significantly differ between the polyandrous *P. maniculatus* (red dots) and its monogamous sister species, *P. polionotus* (blue dots). The calcium concentration between these regions significantly differed only for *P. maniculatus*. Small dots denote values measured for individual females; large dots represent the mean values for each species.

Within both species, the fluid collected from the uterus had a significantly higher pH than the oviduct (paired t-test: *P. maniculatus t* = 21, df = 9, *p* < 0.001; *P. polionotus t* = 6.986, df = 9, *p* < 0.001; Figure 2). In *P. maniculatus*, the fluid collected from the uterus was significantly more calcemic (paired t-test: *t*= 3.95, df = 9, *p* < 0.01), a pattern that was not observed in *P. polionotus* (paired t-test: *t* = 1.98, df = 9, *p*= 0.079).

## DISCUSSION

Reproductive traits are among the most rapidly evolving traits in nature and are driven by many evolutionary processes, including post-copulatory sexual selection (e.g., Eberhard 2004; Clark et al. 2006; Martin-Coello et al. 2009; Ramm et al. 2009). In polyandrous systems, sperm from multiple partners are expected to interact and compete for fertilization of the available ova, and females are predicted to evolve traits that allow them to exert control over the outcome of this competition (reviewed in Eberhard 1996; Firman et al. 2017). In this study, we asked whether female reproductive traits that impact the successful migration of sperm to the site of fertilization differ between polyandrous and monogamous species. We examined the composition of fluids collected from the two reproductive organs closest to the fertilization site – the oviduct and the uterus – using *Peromyscus* mice with naturally variable mating systems among closely related species. Our results show that (1) polyandrous species have significantly more viscous fluid in the uterus but less viscous fluid in the oviduct than monogamous species, and a viscosity gradient from the uterus to the oviduct increases in monogamous species but decreases in polyandrous species; (2) the reproductive fluid pH is significantly higher in the uterus and oviduct of the polyandrous *P. maniculatus* compared to its monogamous congener, *P. polionotus*, but both species have a more alkaline environment in the uterus than oviduct overall; and (3) there are no differences in the calcium content between species, but *P. maniculatus* has a calcium gradient that decreases at the distal end of the reproductive tract. Given that these traits and their interactions are likely to affect sperm motility and migration toward the site of fertilization, these fluidic properties warrant further study to determine the extent to which they provide females control of sperm use within these mice.

Using a highly sensitive method (Duncan et al. 2016), we found that polyandrous species (*P. maniculatus, P. leucopus*, and *P. gossypinus*) have significantly more viscous fluid in the uterus but less viscous fluid in the oviduct than their closely related monogamous congeners (*P. californicus, P. eremicus*, and *P. polionotus*). *In vitro* experiments in *Peromyscus* have shown that increasing the viscosity of the microenvironment leads to reduced sperm velocity (Hook et al. 2022), consistent with other systems (Smith et al. 2009; Miki and Clapham 2013). Our findings suggest that in polyandrous species, sperm motility is hindered more in the uterus, which is closest to the sperm entry site and therefore provides an initially more competitive environment that likely selects for the most motile sperm prior to ever reaching the oviduct (Holt and Fazeli 2016; Suarez 2016). The opposite pattern was observed in the monogamous species, in which sperm motility would be enhanced in the uterus but hindered in the oviduct. In these monogamous species, it is possible that the UTJ – a constricted passageway that separates these two tract regions and contains highly viscous fluid in other species (reviewed in Hunter 1995) – provides an adequately effective barrier to remove morphologically abnormal and slower sperm (Chatdarong et al. 2004; Druart 2012), as has been observed in a previous study examining their collective sperm groups (Hook et al. 2022), and that the higher oviductal fluid viscosity is effective in reducing sperm motility and, thus, the possibility of polyspermy (Kim et al. 1996; Firman and Simmons 2013; Firman 2018). Overall, the significant differences in fluid viscosity based on mating systems is suggestive of an association with post-copulatory sexual selection. Further studies are warranted in these species to examine how sperm interact with uterine and oviductal fluids, as well as seminal fluids after mating (Miki and Clapham 2013), to transverse the UTJ and reach the fertilization site given these observed differences in female fluidic viscosity.

We also found that the reproductive fluid pH is significantly higher in the uterus and oviduct of the polyandrous deer mouse (*P. maniculatus*) during estrus compared to its monogamous congener, *P. polionotus*, but that both species have a more alkaline environment in the uterus than oviduct. This result is surprising given that the opposite has been observed in other studies (López-Albors et al. 2021, but see Ng et al. 2018). Such alkaline environments have been shown to enhance sperm motility in birds (*Gallus domesticus, Coturnix coturnix, Meleagris gallopavo*; Holm and Wishart 1998) and humans (Saito et al. 1996; Zhou et al. 2015), and to activate sperm-specific CatSper channels in mice (*Mus musculus*; Kirichok et al. 2006) and humans (Lishko et al. 2010), thereby inducing sperm hyperactivation (Suarez et al. 1993). Sperm hyperactivation, which takes place in the lower oviduct in rabbits (Overstreet and Cooper 1979) and mice (Suárez and Osman 1987), is an important sperm movement pattern characterized by a deep flagellar bend (reviewed in Suarez and Ho 2003). In bulls, sperm cells exhibit deep asymmetrical bends in pH 7.9-8.5 solutions and shallow asymmetrical bends in pH 7.0-7.5 solutions (Ho et al. 2002). We found that the average pH in upper oviduct fluid was 7.44 in *P. maniculatus* and 7.28 in *P. polionotus*, which is consistent with shallow asymmetrical bends in hyperactivated sperm in both species. Our finding that this trait differs between species based on mating system suggests it might be a trait that is driven by post-copulatory sexual selection, although the greater pH of fluids in the polyandrous species suggests this trait is enabling them to have greater sperm motility, rather than hindering their movement or serving as a barrier for movement. Further studies are warranted to verify the effects of pH on *Peromyscus* sperm movements *in vivo* and how this trait synergistically interacts with the viscosity or calcemic contents of female reproductive fluids or impacts the ability of sperm to capacitate (Stival et al. 2016) or form collective groups that swim together (Fisher and Hoekstra 2010; Hook et al. 2022).

Last, we found that the concentration of calcium in the uterine and oviductal fluids extracted from the polyandrous *P. maniculatus* and monogamous *P. polionotus* did not significantly differ. If this fluidic component enables female control of sperm use, we would expect a difference between these species that differ by mating system. However, it is interesting to note that only in *P. maniculatus* did we observe a gradient in which calcium concentration decreases moving up the female reproductive tract from the uterus to the oviduct. There is a positive association between extracellular calcium and sperm velocity in humans (Zhou et al. 2015), rats (Lindemann and Goltz 1988), and hamsters (Suarez and Dai 1995), which suggests that the reduction in calcemic contents in the oviductal fluids of the polyandrous *P. maniculatus* may impose a barrier on sperm motility. Alternatively, it may indicate that sperm hyperactivate much earlier in *P. maniculatus*, given that calcium is essential for sperm hyperactivation (Yanagimachi 1982), and increases in extracellular calcium levels increase calcium entry through sperm-specific CatSper channels (Marquez and Suarez 2007). We cannot rule out that this property interacts with other fluidic properties or with seminal fluids to control sperm migration, so future studies that examine these effects – specifically through *in vivo* experiments and accounting for potential interspecies variation in these fluidic properties – are warranted.

Our results demonstrate some important differences in the fluid collected from different regions of the female reproductive tract, however finer-scale changes in more localized areas of the reproductive tract and in response to seminal fluids will further enhance our understanding of how selection has shaped the fertilization environment in *Peromyscus*. Our understanding of the physiological mechanisms required for mammalian fertilization remain obscure without the ability to measure conditions *in vivo* in real-time (Ng et al. 2018), especially in small animals, but our results suggest that even closely-related species may exhibit striking differences similar in magnitude to differences in highly divergent taxa (López-Albors et al. 2021). In other mammals, evidence suggests that uterine fluid near the cervix is more viscous than more proximal regions of the uterus, and oviductal fluid in the ampullary region contains viscous compounds produced from ovulating follicles and peritoneal fluid during estrus (reviewed in Hunter et al. 2011). In bovine, for example, the greater amount mucus in the ampulla compared to lower regions of the oviduct is associated with reduced sperm numbers near the fertilization site (Suarez et al. 1997). Our study was limited by the small quantity of fluid we could extract from *Peromyscus* oviducts – although we aimed to maximize this with our approach. The comparisons we were able to make between fluid obtained from the isthmus and ampullary regions suggest that fluidic properties in the oviduct may be equally as dynamic as the structural features (e.g., Yániz et al. 2000; Suarez 2016; Miller 2018).

Taken together, our results support the prediction that female reproductive fluids can vary by the species’ mating system, and thus level of post-copulatory sexual selection, yet the directionality of the differences make their functional significance less clear. We found that female reproductive fluids in polyandrous species is more acidic with differing viscosities throughout the tract compared to monogamous *Peromyscus* females. We also found variation between distinct regions of the reproductive tract, providing indirect evidence for how these properties might impact sperm cells as they migrate up toward the site of fertilization. Our results suggest that fluid viscosity and pH may provide promising avenues for investigating a female reproductive trait that is driven by cryptic female choice, although follow-up experiments are needed to assess their impacts on sperm motility *in vivo* and on male fertilization success within a competitive context.

## Data Availability Statement

All data are available in Dryad.

## Author Contributions

KAH, HSF, and CL conceived of the study, designed experiments, and interpreted results; CL, KAH, and KAJ collected the data, KAJ analyzed video data, KAH carried out the statistical analyses; all authors wrote the manuscript and all authors gave final approval for publication.

## Acknowledgements

We are grateful to Hopi Hoekstra for providing *P. gossypinus* males and to Erica Glasper for providing *P. californicus* males and providing use of a microplate reader. Thanks to W. David Weber for help in determining estrous phase of female mice, maintaining the mouse colony, and collecting many of the reproductive tracts analyzed for this study. We thank Mollie Manier, Halli Weiner, and Patricia Martin-DeLeon for advice on methods for measuring calcium and pH and Shelby Wilson for statistical advice. Thanks to Harrison Arsis, Madeline-Sophie Dang, Catherine Liu, and Audrey Mvemba for their assistance with video analyses. Funding was provided by a National Science Foundation Postdoctoral Research Fellowship [1711817] to KAH, a University of Maryland Honors College Research Grant to CL, the Cystic Fibrosis Foundation (JOYNER18FO) to KAJ, a Burroughs Wellcome Fund Career Award at the Scientific Interface to GAD, and a Eunice Kennedy Shriver National Institute of Child Health and Human Development K99/R00 Pathway to Independence Award [R00HD071972] to HSF.

## LITERATURE CITED

Anderson, M. J., A. S. Dixson, and A. F. Dixson. 2006. Mammalian sperm and oviducts are sexually selected: evidence for co-evolution. Journal of Zoology 270:682–686.

Baker, R. R., and M. A. Bellis. 1993. Human sperm competition: ejaculate manipulation by females and a function for the female orgasm. Animal Behaviour 46:887–909.

Bedford, N. L., and H. E. Hoekstra. 2015. The natural history of model organisms: Peromyscus mice as a model for studying natural variation. eLife 4:e06813.

Brokaw, C. J., R. Josslin, and L. Bobrow. 1974. Calcium ion regulation of flagellar beat symmetry in reactivated sea urchin spermatozoa. Biochemical and Biophysical Research Communications 58:795–800.

Chatdarong, K., C. Lohachit, and C. Linde-Forsberg. 2004. Distribution of spermatozoa in the female reproductive tract of the domestic cat in relation to ovulation induced by natural mating. Theriogenology 62:1027–1041.

Clark, N. L., J. E. Aagaard, and W. J. Swanson. 2006. Evolution of reproductive proteins from animals and plants. Reproduction 131:11–22. Society for Reproduction and Fertility.

Crocker, J. C., and D. G. Grier. 1996. Methods of Digital Video Microscopy for Colloidal Studies. Journal of Colloid and Interface Science 179:298–310.

Druart, X. 2012. Sperm Interaction with the Female Reproductive Tract. Reproduction in Domestic Animals 47:348–352.

Duncan, G. A., J. Jung, A. Joseph, A. L. Thaxton, N. E. West, M. P. Boyle, J. Hanes, and J. S. Suk. 2016. Microstructural alterations of sputum in cystic fibrosis lung disease. JCI Insight 1:e88198.

Eberhard, W. 1996. Female Control: Sexual Selection by Cryptic Female Choice. Princeton, NJ: Princeton University Press.

Eberhard, W. G. 2004. Rapid Divergent evolution of sexual morphology: comparative tests of antagonistic coevolution and traditional female choice. Evolution 58:1947–1970.

Edvardsson, M., and A. Göran. 2000. Copulatory courtship and cryptic female choice in red flour beetles Tribolium castaneum. Proceedings of the Royal Society of London. Series B: Biological Sciences 267:559–563.

Firman, R. C. 2020. Of mice and women: advances in mammalian sperm competition with a focus on the female perspective. Philosophical Transactions of the Royal Society B: Biological Sciences 375:20200082.

Firman, R. C. 2018. Postmating sexual conflict and female control over fertilization during gamete interaction. Annals of the New York Academy of Sciences 1422:48–64.

Firman, R. C., C. Gasparini, M. K. Manier, and T. Pizzari. 2017. Postmating Female Control: 20 Years of Cryptic Female Choice. Trends in Ecology & Evolution 32:368–382.

Firman, R. C., and L. W. Simmons. 2013. Sperm competition risk generates phenotypic plasticity in ovum fertilizability. Proceedings of the Royal Society B: Biological Sciences 280:20132097.

Fisher, H. S., and H. E. Hoekstra. 2010. Competition drives cooperation among closely related sperm of deer mice. Nature 463:801.

Gasparini, C., A. Pilastro, and J. P. Evans. 2020. The role of female reproductive fluid in sperm competition. Philosophical Transactions of the Royal Society B: Biological Sciences 375:20200077.

Gomendio, M., and E. R. S. Roldan. 1993. Coevolution between male ejaculates and female reproductive biology in eutherian mammals. Proceedings of the Royal Society of London. Series B: Biological Sciences 252:7–12.

Hanson, F. W., and J. W. Overstreet. 1981. The interaction of human spermatozoa with cervical mucus in vivo. American Journal of Obstetrics and Gynecology 140:173–178.

Ho, H. C., K. A. Granish, and S. S. Suarez. 2002. Hyperactivated motility of bull sperm is triggered at the axoneme by Ca2+ and not cAMP. Developmental Biology 250:208–217.

Ho, H. C., and S. S. Suarez. 2001. An inositol 1,4,5-trisphosphate receptor-gated intracellular Ca2+ store is involved in regulating sperm hyperactivated motility1. Biology of Reproduction 65:1606–1615.

Holm, L., and Y. Ridderstråale. 1998. Localization of carbonic anhydrase in the sperm-storing regions of the turkey and quail oviduct. The Histochemical Journal 30:481–488.

Holm, L., and G. J. Wishart. 1998. The effect of pH on the motility of spermatozoa from chicken, turkey and quail. Animal Reproduction Science 54:45–54.

Holt, W. V., and A. Fazeli. 2016. Sperm selection in the female mammalian reproductive tract. Focus on the oviduct: Hypotheses, mechanisms, and new opportunities. Theriogenology 85:105–112.

Hook, K. A., W. D. Weber, and H. S. Fisher. 2022. Postcopulatory sexual selection is associated with sperm aggregate quality in Peromyscus mice. Behavioral Ecology 33:55–64.

Hunter, R. H. F. 1995. How, when, and where do spermatozoa gain their fertilising ability in vivo? Reproduction in Domestic Animals 31:51–55.

Hunter, R. H. F., P. Coy, J. Gadea, and D. Rath. 2011. Considerations of viscosity in the preliminaries to mammalian fertilisation. Journal of Assisted Reproduction and Genetics 28:191–197.

Johansson, M., P. Tienthai, and H. Rodríguez-Martínez. 2000. Histochemistry and ultrastructure of the intraluminal mucus in the sperm reservoir of the pig oviduct. Journal of Reproduction and Development 46:183–192.

Joyner, K., D. Song, R. F. Hawkins, R. D. Silcott, and G. A. Duncan. 2019. A rational approach to form disulfide linked mucin hydrogels. Soft Matter 15:9632–9639.

Kim, N.-H., H. Funahashi, L. R. Abeydeera, S. J. Moon, R. S. Prather, and B. N. Day. 1996. Effects of oviductal fluid on sperm penetration and cortical granule exocytosis during fertilization of pig oocytes in vitro. Reproduction 107:79–86.

Kirichok, Y., B. Navarro, and D. E. Clapham. 2006. Whole-cell patch-clamp measurements of spermatozoa reveal an alkaline-activated Ca2+ channel. Nature 439:737–740.

Kirkman-Brown, J. C., and D. J. Smith. 2011. Sperm motility: is viscosity fundamental to progress? Molecular Human Reproduction 17:539–544.

Lenth, R. V. 2016. Least-squares means: The R package “lsmeans.” Journal of Statistical Software 69:1– 33.

Lindemann, C. B., and J. S. Goltz. 1988. Calcium regulation of flagellar curvature and swimming pattern in triton X-100–extracted rat sperm. Cell Motility 10:420–431.

Lishko, P. V., I. L. Botchkina, A. Fedorenko, and Y. Kirichok. 2010. Acid extrusion from human spermatozoa is mediated by flagellar voltage-gated proton channel. Cell 140:327–337.

López-Albors, O., P.J. Llamas-López, J. Á. Ortuño, R. Latorre, and F.A. García-Vázquez. 2021. In vivo measurement of pH and CO2 levels in the uterus of sows through the estrous cycle and after insemination. Sci Rep 11:3194.

Lüpold, S., J. B. Reil, M. K. Manier, V. Zeender, J. M. Belote, and S. Pitnick. 2020. How female × male and male × male interactions influence competitive fertilization in Drosophila melanogaster. Evolution Letters 4:416–429.

Manier, M. K., J. M. Belote, K. S. Berben, D. Novikov, W. T. Stuart, and S. Pitnick. 2010. Resolving mechanisms of competitive fertilization success in Drosophila melanogaster. Science 328:354– 357.

Marquez, B., and S. S. Suarez. 2007. Bovine sperm hyperactivation is promoted by alkaline-stimulated Ca2+ influx. Biology of Reproduction 76:660–665.

Martin-Coello, J., H. Dopazo, L. Arbiza, J. Ausió, E. R. S. Roldan, and M. Gomendio. 2009. Sexual selection drives weak positive selection in protamine genes and high promoter divergence, enhancing sperm competitiveness. Proceedings of the Royal Society B: Biological Sciences 276:2427–2436.

Miki, K., and D. E. Clapham. 2013. Rheotaxis guides mammalian sperm. Current Biology 23:443–452.

Miller, D. J. 2018. Review: The epic journey of sperm through the female reproductive tract. Animal 12:s110–s120.

Ng, K. Y. B., R. Mingels, H. Morgan, N. Macklon, and Y. Cheong. 2018. In vivo oxygen, temperature and pH dynamics in the female reproductive tract and their importance in human conception: a systematic review. Human Reproduction Update 24:15–34.

Orr, T. J., M. Burns, K. Hawkes, K. E. Holekamp, K. A. Hook, C. C. Josefson, A. A. Kimmitt, A. K. Lewis, S. E. Lipshutz, K. S. Lynch, L. K. Sirot, D. J. Stadtmauer, N. L. Staub, M. F. Wolfner, and V. Hayssen. 2020. It takes two to tango: including a female perspective in reproductive biology. Integrative and Comparative Biology 60:796–813.

Overstreet, J. W., and G. W. Cooper. 1979. Effect of ovulation and sperm motility on the migration of rabbit spermatozoa to the site of fertilization. Reproduction 55:53–59.

Parker, G. A. 1970. Sperm competition and its evolutionary consequences in the insects. Biological Reviews 45:525–567.

Patczai, B., T. Mintál, L. G. Nőt, N. Wiegand, and D. Lőrinczy. 2017. Effects of deep-freezing and storage time on human femoral cartilage. Journal of Thermal Analysis and Calorimetry 127:1177–1180.

Petrunkina, A. M., J. Friedrich, W. Drommer, G. Bicker, D. Waberski, and E. Töpfer-Petersen. 2001. Kinetic characterization of the changes in protein tyrosine phosphorylation of membranes, cytosolic Ca2+ concentration and viability in boar sperm populations selected by binding to oviductal epithelial cells. Reproduction 122:469–480.

Pizzari, T., and T. R. Birkhead. 2000. Female feral fowl eject sperm of subdominant males. Nature 405:787–789.

R Core Team. 2016. R: A Language and Environment for Statistical Computing. Vienna, Austria. R Foundation for Statistical Computing. Retrieved from https://www.R-project.org/

Ramm, S. A., L. McDonald, J. L. Hurst, R. J. Beynon, and P. Stockley. 2009. Comparative proteomics reveals evidence for evolutionary diversification of rodent seminal fluid and its functional significance in sperm competition. Molecular Biology and Evolution 26:189–198.

Rodríguez-Martínez, H., F. Saravia, M. Wallgren, P. Tienthai, A. Johannisson, J. M. Vázquez, E. Martínez, J. Roca, L. Sanz, and J. J. Calvete. 2005. Boar spermatozoa in the oviduct. Theriogenology 63:514–535.

Rowe, M., L. Veerus, P. Trosvik, A. Buckling, and T. Pizzari. 2020. The reproductive microbiome: An emerging driver of sexual selection, sexual conflict, mating systems, and reproductive isolation. Trends in Ecology & Evolution 35:220–234.

Saito, K., Y. Kinoshita, H. Kanno, and A. Iwasaki. 1996. The role of potassium ion and extracellular alkalization in reinitiation of human spermatozoa preserved in electrolyte-free solution at 4°C. Fertility and Sterility 65:1214–1218.

Simmons, L. W., and J. L. Fitzpatrick. 2012. Sperm wars and the evolution of male fertility. Reproduction 144:519–534.

Simmons, L. W., and N. Wedell. 2020. Fifty years of sperm competition: the structure of a scientific revolution. Philosophical Transactions of the Royal Society B: Biological Sciences 375:20200060.

Simons, J. E., and S. D. Olson. 2018. Sperm motility: models for dynamic behavior in complex environments. In M. Stolarska and N. Tarfulea (Eds.), Cell Movement (pp. 169–209). Switzerland: Birkhäuser.

Smith, D. J., E. A. Gaffney, H. Gadêlha, N. Kapur, and J. C. Kirkman-Brown. 2009. Bend propagation in the flagella of migrating human sperm, and its modulation by viscosity. Cell Motility and the Cytoskeleton 66:220–236.

Stival, C., L. del C. Puga Molina, B. Paudel, M. G. Buffone, P. E. Visconti, and D. Krapf. 2016. Sperm capacitation and acrosome reaction in mammalian sperm. In M. G. Buffone (Ed.), Sperm Acrosome Biogenesis and Function During Fertilization (pp. 93–106). Switzerland: Springer.

Suarez, S. S. 2008. Control of hyperactivation in sperm. Human Reproduction Update 14:647–657.

Suarez, S. S. 2016. Mammalian sperm interactions with the female reproductive tract. Cell and Tissue Research 363:185–194.

Suarez, S. S., K. Brockman, and R. Lefebvre. 1997. Distribution of mucus and sperm in bovine oviducts after artificial insemination: The physical environment of the oviductal sperm reservoir. Biology of Reproduction 56:447–453.

Suarez, S. S., and X. Dai. 1995. Intracellular calcium reaches different levels of elevation in hyperactivated and acrosome-reacted hamster sperm. Molecular Reproduction and Development 42:325–333.

Suarez, S. S., and H. C. Ho. 2003. Hyperactivation of mammalian sperm. Cellular and Molecular Biology 49:351–356.

Suárez, S. S., and R. A. Osman. 1987. Initiation of hyperactivated flagellar bending in mouse sperm within the female reproductive tract. Biology of Reproduction 36:1191–1198.

Suarez, S. S., S. M. Varosi, and D. Xiaobing. 1993. Intracellular calcium increases with hyperactivation in intact, moving hamster sperm and oscillates with the flagellar beat cycle. Proceedings of the National Academy of Sciences 90:4660–4664.

Thornhill, R. 1983. Cryptic female choice and its implications in the scorpionfly Harpobittacus nigriceps. The American Naturalist 122:765–788.

Troisi, A., and M. Carosi. 1998. Female orgasm rate increases with male dominance in Japanese macaques. Animal Behaviour 56:1261–1266.

Tung, C., L. Hu, A. G. Fiore, F. Ardon, D. G. Hickman, R. O. Gilbert, S. S. Suarez, and M. Wu. 2015. Microgrooves and fluid flows provide preferential passageways for sperm over pathogen Tritrichomonas foetus. Proceedings of the National Academy of Sciences 112:5431–5436.

Tung, C.-K., and S. S. Suarez. 2021. Co-adaptation of physical attributes of the mammalian female reproductive tract and sperm to facilitate fertilization. Cells 10:1297.

Turner, L. M., A. R. Young, H. Römpler, T. Schöneberg, S. M. Phelps, and H. E. Hoekstra. 2010. Monogamy evolves through multiple mechanisms: evidence from V1aR in deer mice. Molecular Biology and Evolution 27:1269–1278.

Vahed, K., and D. J. Parker. 2012. The evolution of large testes: sperm competition or male mating rate? Ethology 118:107–117.

Wickham, H. 2016. ggplot2: Elegant Graphics for Data Analysis. Springer.

Yanagimachi, R. 1982. Requirement of extracellular calcium ions for various stages of fertilization and fertilization-related phenomena in the hamster. Gamete Research 5:323–344.

Yániz, J. L., F. Lopez-Gatius, P. Santolaria, and K. J. Mullins. 2000. Study of the functional anatomy of bovine oviductal mucosa. The Anatomical Record 260:268–278.

Yeung, C.-H., S. Breton, I. Setiawan, Y. Xu, F. Lang, and T. G. Cooper. 2004. Increased luminal pH in the epididymis of infertile c-ros knockout mice and the expression of sodium–hydrogen exchangers and vacuolar proton pump H+-ATPase. Molecular Reproduction and Development 68:159–168.

Yuana, Y., A. N. Böing, A. E. Grootemaat, E. van der Pol, C. M. Hau, P. Cizmar, E. Buhr, A. Sturk, and R. Nieuwland. 2015. Handling and storage of human body fluids for analysis of extracellular vesicles. Journal of Extracellular Vesicles 4:29260.

Zhou, J., L. Chen, J. Li, H. Li, Z. Hong, M. Xie, S. Chen, and B. Yao. 2015. The semen pH affects sperm motility and capacitation. PLOS ONE 10:e0132974.

